# Rational Protein Engineering of Bacterial *N*-demethylases to Create Biocatalysts for the Production of Methylxanthines

**DOI:** 10.1101/2021.12.17.472166

**Authors:** Shelby Brooks Mills, Meredith B. Mock, Ryan M. Summers

## Abstract

Methylxanthines have a rich history as therapeutics and pharmaceuticals. However, natural dimethyl- and monomethylxanthines are difficult to produce synthetically, which has limited further exploration of these compounds in medicinal applications. A biosynthetic method for production of methylxanthines from whole cell biocatalysts is an attractive alternative. The bacterium *Pseudomonas putida* CBB5 contains a set of five enzymes, NdmABCDE, which are responsible for methylxanthine metabolism *via N*-demethylation to xanthine. The recent elucidation of the crystal structures of NdmA and NdmB, which remove the *N*_1_- and *N*_3_-methyl groups of caffeine, respectively, has opened new avenues to create biocatalysts for methylxanthine production. We have created a set of fifteen *N*-demethylase mutants and expressed them in *E. coli* BL21(DE3) as whole cell biocatalysts. The activity of each mutant was characterized for their affinity towards caffeine, theobromine, and theophylline. Two mutant enzymes in particular, labeled NdmA3 and NdmA4, both exhibited selectivity towards the *N*_3_-methyl group instead of the *N*_1_-methyl group. We also discovered that specific point mutations in NdmD resulted in the ability to tune the rate of the *N*-demethylase reaction. These new enzymes provide the capability of producing high-value methylxanthines, such as paraxanthine and 1-methylxanthine, through a biocatalytic route.

## Introduction

Methylxanthines have been used in the pharmaceutical industry for over 100 years, beginning with the use of theophylline as a diuretic and later as an asthma treatment (Barnes 2013). Since then, the use of caffeine (1,3,7-trimethylxanthine), theophylline (1,3-dimethylxanthine), and theobromine (3,7-dimethylxanthine) has only grown more common (Monteiro et al. 2018; Singh et al. 2018; Franco et al. 2013). The majority of therapeutic applications for these methylxanthines have been to target the central nervous, cardiovascular, and respiratory systems, and to serve as smooth muscle relaxants (Ponte et al. 2018; Franco et al. 2013; Singh et al. 2018; Monteiro et al. 2018; Mazzafera 2002). Studies have determined that a habitual intake of caffeine and related methylxanthines can lead to a lower risk of developing Alzheimer’s disease (Eskelinen et al. 2009; Van Gelder et al. 2007; Maia and De Mendonça 2002; Lindsay et al. 2004), depression (Smith 2009; Grosso et al. 2016), and stroke (Larsson 2014; Liebeskind et al. 2016), among other benefits. The low toxicity and important biological effects of methylxanthines make them an ideal group of candidates for therapeutics (Monteiro et al. 2018).

One methylxanthine with interesting therapeutic applications is paraxanthine (1,7-dimethylxanthine). Research has shown that paraxanthine may reduce the risk of developing Parkinson’s disease by protecting nigrostriatal dopaminergic neurons (Guerreiro et al. 2008), acting as a stimulant for the central nervous system (Okuro et al. 2010), and aiding in the treatment of human liver fibrosis (Gressner et al. 2009; Gressner 2009). In rodents, Okuro *et al*. demonstrated that paraxanthine contained lower toxicity and did not increase anxiety levels when compared with caffeine (Okuro et al. 2010). However, despite these benefits, a major challenge for the implementation of paraxanthine as a therapeutic is that there are few viable industrial-scale production options, which results in a high cost of paraxanthine.

There are limited natural resources available to generate and harvest paraxanthine because it is not produced at significant quantities in plants (Camandola et al. 2018; Monteiro et al. 2018). Naturally, paraxanthine is found as a caffeine metabolite in humans, as approximately 84% of ingested caffeine is converted into paraxanthine before being metabolized further (Guerreiro et al. 2008; Costenla et al. 2010; Costenla et al. 1999). The current chemically synthetic process employed to generate methylxanthines is not optimal because selective alkylation of the nitrogen atoms is difficult to control, toxic, and costly (Dash and Gummadi 2006; Gopishetty 2012). In place of a synthetic chemistry procedure, we are interested in developing a biocatalytic to produce paraxanthine method through biotransformation of caffeine. Biotransformations have numerous advantages in that they are cost-effective, less energy demanding, eco-friendly, and non-hazardous (Monteiro et al. 2018), while also providing improved selectivity with operation at ambient temperatures (Bornscheuer and Pohl 2001).

The bacterium *Pseudomonas putida* CBB5 metabolizes caffeine via *N*-demethylation using a set of five enzymes, NdmABCDE, belonging to the Rieske non-heme iron monooxygenase family (Yu et al. 2009; Summers et al. 2012; Summers et al. 2013). Caffeine is first *N*_1_-demethylated to theobromine by NdmA, followed by *N*_3_-demethylation to 7-methylxanthine by NdmB (Summers et al. 2012; Algharrawi and Subramanian 2020), although paraxanthine is transiently produced as a minor metabolite (Yu et al. 2009) (Figure 1). Kinetic data of these enzymes confirmed that although theobromine is the preferred substrate for NdmB, the presence of the *N*_1_*-*methyl group on caffeine or theophylline significantly impedes the *N*_3_-demethylation activity of NdmB toward these molecules (Summers et al. 2012). Curiously, while NdmA prefers to remove the *N*_1_-methyl group, the enzyme exhibits a slight promiscuity and is capable of removing the *N*_3_-methyl group from caffeine and theophylline (Algharrawi et al. 2017; Algharrawi et al. 2015). In CBB5, generation of minor amounts of paraxanthine from caffeine was initially attributed to NdmB, but we have observed that this activity is due to promiscuity of NdmA when expressed in *E. coli* (Summers et al. 2012).

**Figure 1.**
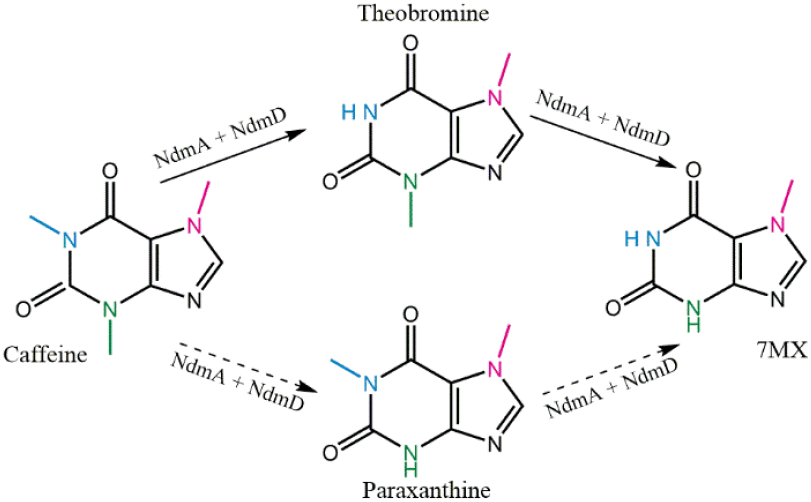
The *N*-demethylation pathway of caffeine to 7-methylxanthine (7MX) performed by NdmA and NdmB with the partnering reductase NdmD. Continuous lines show the preferred pathway while the dashed lines indicate an alternative route.

Recent crystallographic data has identified seven residues in the NdmA active site involved with substrate binding (Kim et al. 2019). When comparing those residues to NdmB through sequence alignment, all but two of the side chain interactions were conserved. The interaction of the NdmD reductase with NdmA was also investigated, and the mutation of the valine 541 residue on NdmD to arginine or tryptophan was shown to reduce *N*-demethylase activity of NdmA, indicating that the electron transfer pathway had been disrupted (Kim et al. 2019). This understanding of protein structures enables rational protein engineering to change the function of a protein through specific mutations (Bornscheuer and Pohl 2001; Harris and Craik 1998).

In this work, we have targeted the two non-conserved residues identified in the NdmA and NdmB crystal structures, creating single and double mutants of each enzyme. These enzymes were expressed in *E. coli* BL21(DE3) and whole cell biocatalysts were characterized to determine the effect of the mutations on enzyme activity toward the methylxanthines caffeine, theophylline, and theobromine. The specificity of the NdmA N282Q F286L double mutant was swapped from the *N*_1_- to the *N*_3_-methyl group. However, mutations to NdmB only served to reduce enzyme activity. We further demonstrated that swapping a loop region from NdmB to the NdmA double mutant greatly increased the rate of paraxanthine formation from caffeine. We have also created *E. coli* strains harboring NdmD mutants to vary the rate of activity by altering the interaction between the reductase and *N*-demethylase. By targeting the V541 residue and removing the N-terminal Rieske cluster on NdmD, we generated a set of strains with varying levels of *N*-demethylase activity controlled at the reductase level. These results expand our capability for biocatalytic production of several methylxanthines, especially the high-value compound paraxanthine.

## Materials and Methods

### Chemicals

All enzymes (Taq, T4 DNA ligase, Phusion) with corresponding buffers were purchased from New England Biolabs (Ipswich, MA). Tryptone, agar, yeast extract, ferric chloride, potassium phosphate dibasic anhydrous, and potassium phosphate monobasic anhydrous were obtained from VWR International (Radnor, PA). Sodium chloride, glacial acetic acid, and HPLC grade methanol were supplied by VWR Chemicals BDH (Radnor, PA). All DNA purification kits were from Omega Bio-Tek (Norcross, GA). The Pfu enzyme was from Agilent (Santa Clara, CA). Caffeine was from J. T. Baker, Avantor (Randor, PA), theophylline was supplied by MP Biomedicals LLC (Irvine, CA), and theobromine was from ACROS organics (Morris Plains, NJ). Isopropyl-β-D-thiogalactopyranoside was purchased from Indofine Chemical company (Hillsborough, NJ). The PCR primers were bought from Eurofins Genomics (Louisville, KY), which also provided DNA sequencing service.

### Plasmid Construction

All plasmids and primers used in this project are listed in Tables S1 and S2, respectively. The NdmA, NdmB, and NdmD mutants were generated by site-directed mutagenesis (Agilent 2015). The A-N282Q-F/R and A-F286L-F/R primer pairs were used with plasmid dDA (Figure S1) to construct plasmids dDA1 and dDA2, respectively. Similarly, plasmids dDB1 and dDB2 were created using primer pairs B-Q289N-F/R and B-L293Q-F/R, respectively, from plasmid dDB (Figure S1). Plasmid dA3 was generated using dDA2 as template with A3-N282A-F/R primers, and dDB2 was mutated to dDB3 using the B3-Q298N-F/R primer set. The V541R and V541W mutations on NdmD were created first using primer pairs D-V541W-F/R and D-V541R-F/R and pET28-His-ndmD (pD) as a template to create plasmids pDW and pDR, respectively. A single point mutation, C69A, was then created from these plasmids using primer pair D-C69A-F/R, resulting in plasmids pD1, pDW1, and pDR1. A second round of mutagenesis to introduce a C50A mutation was carried out with the primer pair D-C50A-F/R, generating plasmids pD2, pDW2 and pDR2. A loop-swapped NdmA_QL_ mutant (Kim et al. 2019) was amplified using primers Loop-F-NdeI and NdmA-R-KpnI and cloned into the pACYCDuet-1 plasmid already containing the *ndmD* gene, generating plasmid dDA4. Successful mutagenesis and cloning was confirmed by DNA sequencing. All plasmids were transformed into *E. coli* BL21(DE3) for protein production using a modified heat shock method based on Chung *et al*. (Chung et al. 1989). A full list of strains used in this project is given in Table S3.

### Cell Growth and Assays

Induction of gene expression and resting cell assays were carried out as described previously (Yu et al. 2009; Quandt et al. 2013; Summers et al. 2013). Cells containing the plasmids of interest were grown in LB broth with appropriate antibiotics at 37°C with 200 rpm shaking. When the OD_600_ reached 0.5, FeCl_3_ was added to a final concentration of 10 μM, and the cultures were moved to 18°C with 200 rpm shaking. At an OD_600_ of 0.8-1.0, IPTG was added to a final concentration of 0.1 mM, and the cultures were grown overnight at 18°C for 16-20 h. Cells were harvested by centrifugation at 10,000 × g for 10 minutes at 4°C and resuspended in 50 mM potassium phosphate (KP_i_) buffer (pH 7.5).

Resting cell assays were carried out in a 2 mL total volume reaction containing 1 mM methylxanthine and freshly-harvested whole cells (OD_600_ = 5.0) in KP_i_ buffer. Reactions were incubated at 30°C with 200 rpm agitation. A sample from each reaction was taken at various time points and combined with an equal amount of acetonitrile or methanol to stop the reaction from proceeding. All samples were performed in triplicate.

### Analytical Procedures

Samples from resting cell assays were analyzed with a Shimadzu LC-20AT high performance liquid chromatography (HPLC) equipped with an SPD-M30A photodiode array detector to identify metabolic products and quantify the methylxanthines as described previously (Summers et al. 2012; Yu et al. 2009). Compounds were separated on a Hypersil BDS C_18_ column (100 mm × 4.6 mm) with a mobile phase of methanol/water/acetic acid (15:85:0.5 /v/v).

## Results

Recent crystal structures of NdmA and NdmB revealed two domains: a Rieske domain at the N-terminus containing 3 α-helices and 9 β-strands and a C-terminal ligand binding domain with 5 α-helices and 8 β-strands (Kim et al. 2019). Additionally, both possess a loop region between β-13 and β-14 that is highly flexible and changes conformation depending upon the substrate binding. Within the binding pocket of NdmA, F168 established a π-π bond with the purine backbone of the caffeine molecule, with additional hydrophobic interactions between caffeine and side chain atoms of F174, F223, L248, N282, V285, and F286. These residues allow alignment of caffeine in the binding pocket to enable removal of the *N*_1_-methyl group. Interestingly, the sequence alignment indicated that all but two of the same residues were present in the NdmB binding pocket, which is responsible for *N*_3_-demethylation of theobromine; the N282 and F286 in NdmA are Q289 and L293 in NdmB. Indeed, an NdmA N282Q F286L double mutant (previously named NdmA_QL_, but hereafter termed NdmA3) demonstrated increased activity toward the *N*_3_-methyl group of caffeine in purified enzyme studies (Kim et al. 2019). As enzyme activity can vary greatly when assayed *in vivo* compared with *in vitro* reactions, we sought to determine the activity of mutated NdmA and NdmB enzymes in whole cells.

Single mutants of NdmA N282Q (NdmA1), F286L (NdmA2), NdmB Q289N (NdmB1), NdmB L293F (NdmB2), and double mutants NdmA3 and NdmB Q289N L293F (NdmB3) were constructed through site-directed mutagenesis. Each gene was placed in the pACYCDuet-1 plasmid containing the *ndmD* gene in a separate multiple cloning site. The NdmD enzyme is a unique Rieske reductase that is absolutely essential for *N*-demethylation activity, passing electrons from NADH to NdmA or NdmB (Summers et al. 2013). Each *N*-demethylase mutant-reductase combination was expressed in *E. coli* BL21(DE3), and activity of whole cells toward caffeine, theobromine, and theophylline was determined through resting cell assays.

### Activity of NdmA mutants

Each NdmA mutation decreased the activity of the enzyme toward caffeine (Figure 2A, Table 1). The F286L single mutation had the lowest effect on enzyme activity, with reduction in caffeine consumption rate about 1.5-fold in cells expressing *ndmA2* compared with the controls expressing the wild-type *ndmA*. The largest change in activity was due to the N282Q mutation, which reduced activity in cells with NdmA1 by over 15-fold. Surprisingly, the NdmA3 double mutant improved cell activity over that of NdmA1 by 2.7-fold, with only a 7.2-fold reduction in activity from the wild-type NdmA. This reduction in activity is considerably less than the 18-fold reduction in activity observed by the double mutant in purified enzyme assays (Kim et al. 2019).

**Table 1.** Comparison of methylxanthine consumption for NdmA and each NdmA mutant after two hours

| Strain | Caffeine Consumed ($\mu\text{M}$) | Rate of Caffeine Consumption ($\mu\text{M}/\text{minute}$) | Percent of Paraxanthine in Products* | Theophylline Consumed ($\mu\text{M}$) | Rate of Theophylline Consumption ($\mu\text{M}/\text{minute}$) | Percent of 1-Methylxanthine in Products** |
| --- | --- | --- | --- | --- | --- | --- |
| dDA | $1109.7 \pm 21.9$ | $10.3 \pm 1.2$ | $1.4 \pm 0.5$ | $830.2 \pm 16.1$ | $6.7 \pm 0.1$ | $7.4 \pm 0.2$ |
| dDA1 | $96.4 \pm 78.1$ | $0.5 \pm 0.5$ | $28.9 \pm 5.3$ | $78.9 \pm 26.0$ | $0.8 \pm 0.3$ | $88.1 \pm 0.1$ |
| dDA2 | $850.5 \pm 39.8$ | $6.9 \pm 0.1$ | $5.1 \pm 0.3$ | $810.4 \pm 51.6$ | $6.7 \pm 0.4$ | $17.0 \pm 0.1$ |
| dDA3 | $185.8 \pm 35.1$ | $1.4 \pm 0.2$ | $82.9 \pm 0.5$ | $784.0 \pm 20.6$ | $6.4 \pm 0.2$ | $98.6 \pm 0.2$ |

\* Percent of the consumed caffeine converted to paraxanthine instead of theobromine
\*\* Percent of the consumed theophylline converted to 1-methylxanthine instead of 3-methylxanthine

**Figure 2.**
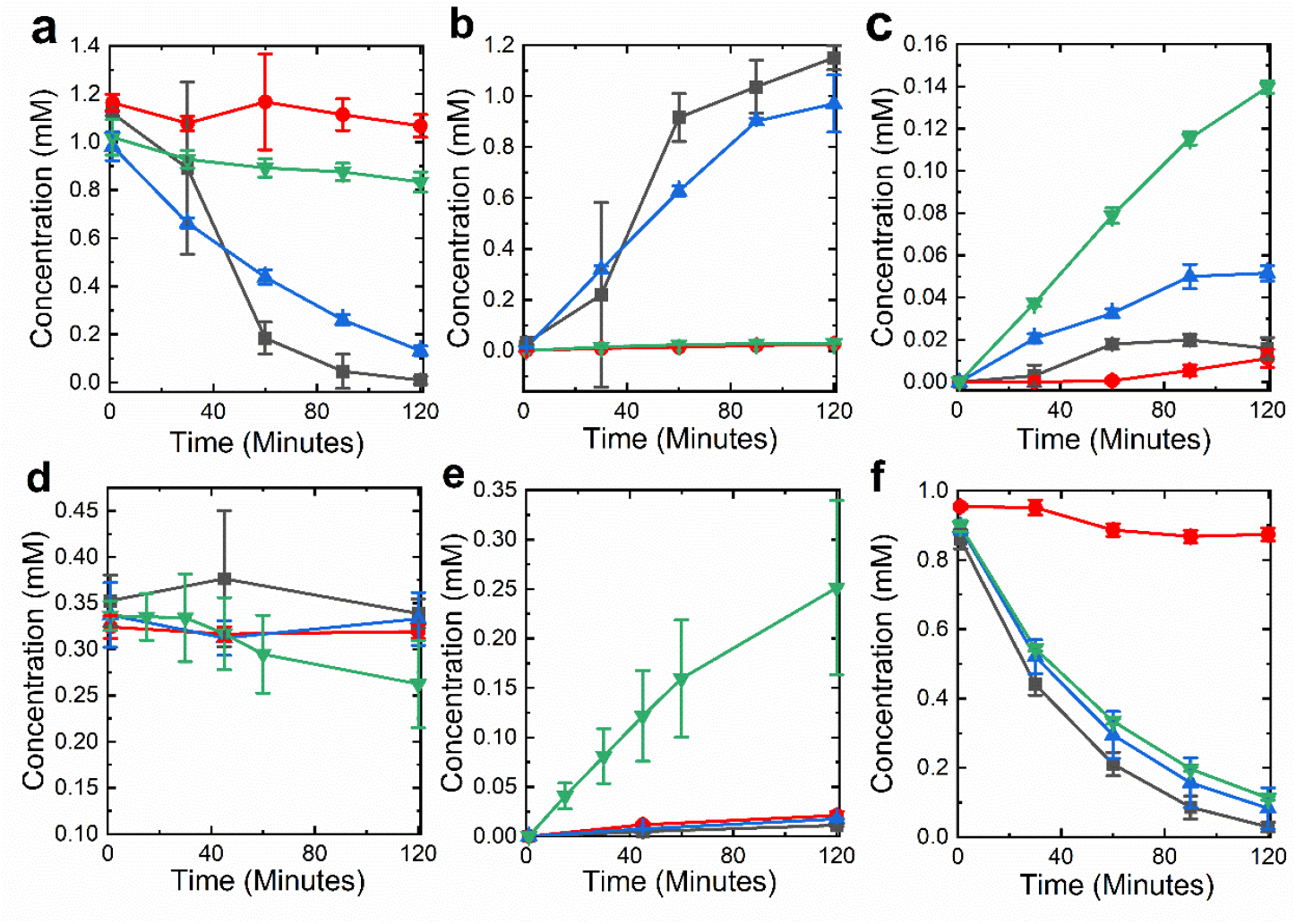
Whole cells containing NdmA, NdmA1, NdmA2, and NdmA3 are active toward several methylxanthines. Caffeine (a) is converted to theobromine (b) and paraxanthine (c), and theobromine (d) to 7-methylxanthine (e). (f) Degradation of theophylline. ■, NdmA control; ●, NdmA1; ▲, NdmA2; ▼, NdmA3. Cells (OD_600_ = 5.0) were incubated with 1 mM caffeine or theophylline or 0.35 mM theobromine in 50 mM KP_i_ buffer at 30°C with 200 rpm shaking, and metabolites were quantified by HPLC. Concentrations reported are means with standard deviations of triplicate results.

A closer look at the metabolites produced from caffeine (Figure 2B & C) demonstrates that the N282 residue is key in aligning caffeine in the NdmA binding pocket for *N*_1_-demethylation. The N282Q mutation increased the amount of paraxanthine in the products from caffeine from 1.4% by the wild-type NdmA to 28.9% by NdmA1, whereas the single F286L mutant NdmA2 only produced 5.1% paraxanthine (Table 1). Furthermore, the double mutant displayed high synergy in shifting activity from the *N*_1_- to the *N*_3_-methyl group, resulting in 82.9% paraxanthine produced from the caffeine consumed. When theobromine was used as substrate to test activity toward the *N*_3_-methyl group, cells containing NdmA, NdmA1, and NdmA2 showed no significant decrease in theobromine concentration (Figure 2D). However, cells with NdmA3 exhibited slight *N*_3_-demethylase activity toward theobromine, consuming 74.0 ± 32.7 µM theobromine over two hours (Figure 2D & E). Thus, the N282Q and F286L mutations enable the NdmA3 binding pocket to mimic that of NdmB, although at lower positional specificity and kinetic rate.

Replacing caffeine with theophylline in the resting cell assay further confirmed the above results. The N282Q mutation (NdmA1) resulted in 88.1 % of the marginal theophylline consumed being converted to 1-methylxanthine instead of 3-methylxanthine (Figure 2F, Table 1). The addition of the second mutation, F286L (NdmA3) further increased the amount of 1-methylxanthine as product to 98.6% and increased the enzymatic activity when compared to NdmA1 (Table 1). Strain dDA3 expressing the double mutant reacted 4.5 times faster towards theophylline than caffeine, indicating that theophylline may fit better in the NdmA3 binding pocket due to the lack of an *N*_7_-methyl group.

### Activity of NdmB mutants

We began characterization of the NdmB mutants with theobromine, as it is the preferred substrate for this enzyme (Summers et al. 2012). The control strain expressing *ndmB* with *ndmD* consumed 974.2 ± 19.4 μM theobromine within 90 minutes (Figure 3A). Single mutations resulted in decreased activities toward theobromine (Figure 3A); NdmB1 and NdmB2 cells consumed 793.4 ± 23.1 and 439.8 ±7.0 μM theobromine over two hours. Initial rates of cells containing NdmB1 and NdmB2 toward theobromine were reduced approximately 10-20%, although NdmB1 cells consumed nearly twice as much theobromine as NdmB2 cells over two hours (Figure 3A). Thus, the Q289N mutation on NdmB had a much lower effect on activity than did the N282Q mutation on NdmA, while the effect of the L293F mutation to NdmB was much greater than that of the F286L mutation on NdmA. Cells containing the NdmB3 double mutant exhibit minimal activity toward theobromine, with production of only 40.4 ± 1.3 μM 7-methylxanthine over two hours (Figure 3A & B). This lack of activity toward theobromine by NdmB3 indicated that the binding pocket may be more suitable for *N*_1_-demethylation, as hypothesized.

**Figure 3.**
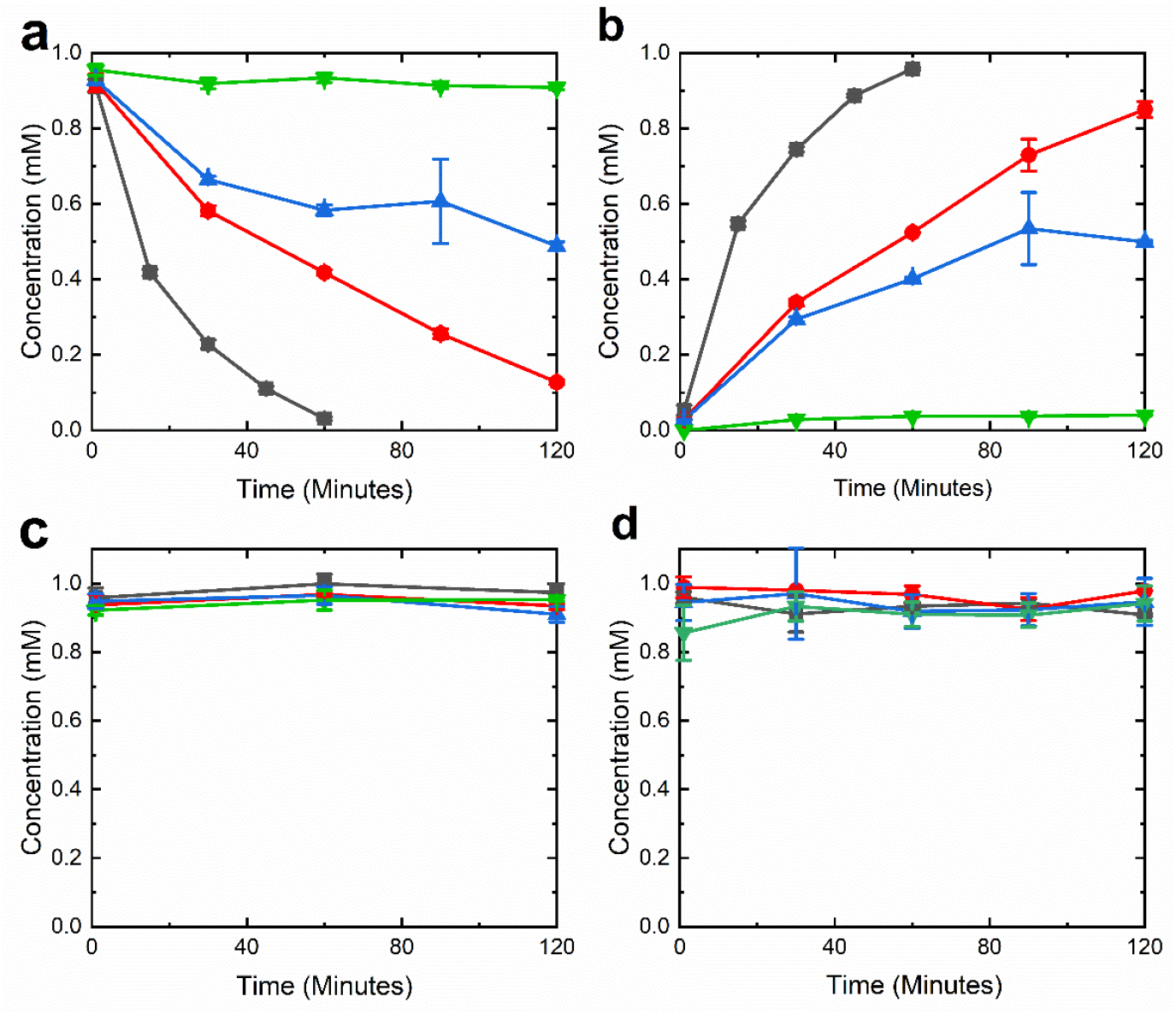
Whole cells containing NdmB and its mutants are only active toward theobromine. Theobromine (a) was converted to 7-methylxanthine (b) at differing rates. No significant activity was observed toward caffeine (c) or theophylline (d). ■, NdmB; ●, NdmB1; ▲, NdmB2; ▼, NdmB3. Concentrations reported are means with standard deviations of triplicate results.

In order to determine whether specificity of NdmB3 was swapped from the *N*_3_- to the *N*_1_-methyl group, we tested activity of the enzyme toward caffeine and theophylline, which both contain *N*_1_-methyl groups. We did not detect a significant change in caffeine (Figure 3C) or theophylline (Figure 3D) concentrations after two hours of incubation with the whole cell biocatalysts. Similarly, consumption of caffeine and theophylline by the single mutants was greatly limited. SDS-PAGE analysis of the cells indicated that NdmB3 was produceded at similar levels as the other mutants (data not shown), thus insolubility of the enzyme is not likely the main reason for lack of activity toward caffeine or theophylline. The wild-type NdmB does not readily *N*_3_-demethylate caffeine or theophylline *in vitro* (Summers et al. 2012), and the same holds true for our resting cell assays. This further suggests that presence the *N*_1_-methyl group inhibits activity of NdmB, most likely through steric hindrances near the active site. Thus, NdmB and the mutants described here are not suitable to perform *N*_1_-demethylation reactions.

### Increased production of paraxanthine

Further analysis of the NdmA crystal structure revealed that a loop region between β13 and β14 changed conformation in the presence of caffeine, and that swapping this loop on the double mutant with the analogous NdmB loop resulted in increased *N*_3_-demethylase activity, albeit at lower protein solubility (Kim et al. 2019). We used the same expression vector described earlier in this study to express the loop-swapped double mutant (hereafter termed NdmA4) in *E. coli* BL21(DE3) cells for direct comparison with the other mutants. Activity of cells containing NdmA4 was similar to those of cells with NdmA3 when caffeine was used as substrate (Figure 4A & B), although molar yield of paraxanthine was higher with NdmA4 (Table 2). We also observed slight production of 7-methylxanthine, generated by *N*_1_-demethylation of paraxanthine (data not shown).

**Table 2.** Comparison of paraxanthine production over two hours by whole cells containing NdmA, NdmA3, and NdmA4.

| Strain | Caffeine Consumed ($\mu\text{M}$) | Paraxanthine in Products ($\mu\text{M}$) | Molar Yield |
| --- | --- | --- | --- |
| dDA | $1109.7 \pm 21.9$ | $15.9 \pm 5.0$ | $0.01 \pm 0.003$ |
| dDA3 | $105.0 \pm 16.1$ | $92.4 \pm 3.1$ | $0.77 \pm 0.11$ |
| dDA4 | $88.6 \pm 32.5$ | $83.9 \pm 3.7$ | $0.97 \pm 0.32$ |

**Figure 4.**
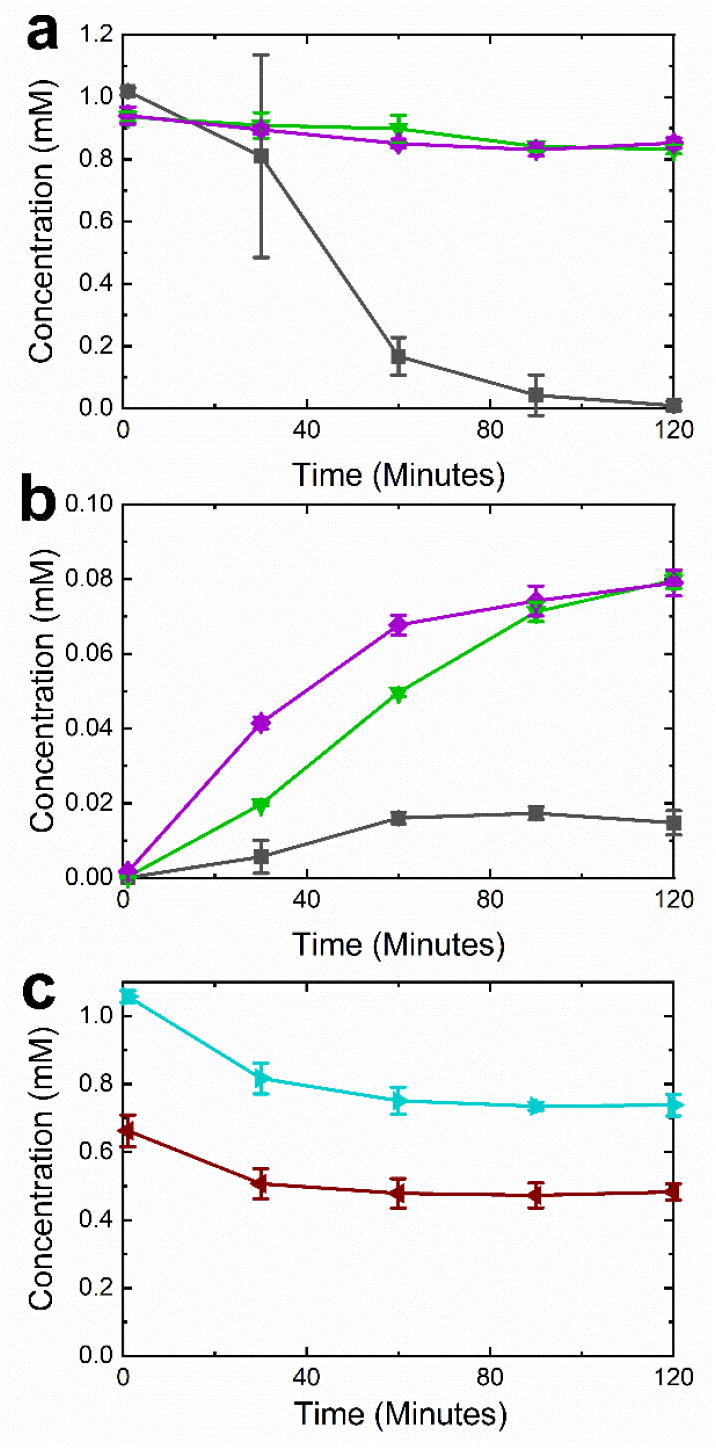
Cells containing NdmA4 are also active toward various methylxanthines. Caffeine (a) is converted to paraxanthine (b) over the course of two hours by cells containing NdmA (■), NdmA3 (▼), and NdmA4 (♦). (c) NdmA4 cells are also capable of consuming theophylline (►) and theobromine (◀). Concentrations reported are means with standard deviations of triplicate results.

Cells containing NdmA4 had decreased activity and conversion of theophylline compared to cells with NdmA3. Over the same two hour period only 30% of the theophylline was consumed by cells with NdmA4 (Figure 4C) in contrast to the 87% of theophylline consumption achieved by cells expressing NdmA3 (Figure 2F). Increased *N*_3_-demethylase activity with NdmA4 cells was more readily detected when theobromine was used as substrate (Figure 4C). Cells containing NdmA4 consumed 180 ± 32 μM theobromine in 120 min, compared with 74 ± 33 μM theobromine consumed by cells with NdmA3 (Figure 2D) over the same time period. This further confirms the mutations on NdmA to mimic the environment of the NdmB binding pocket were successful, and this protein engineering approach switched the selectivity from the *N*_1_-methyl group to the *N*_3_-methyl group of caffeine and related methylxanthines.

### NdmD mutants enable additional tuning of activity

Initial reports using purified enzymes suggested that mutating the V541 residue of NdmD could interrupt the electron transfer from the reductase to the *N*-demethylase; a V541W mutation (NdmDW) reduced *N*-demethylation activity of NdmA to below 20% that of the wild type while a V541R mutation (NdmDR) resulted in almost no activity (Kim et al. 2019). The differences in *N*-demethylase activity between *in vitro* and *in vivo* conditions demonstrated in this study led us to investigate whether the reductase mutants would also act differently in our whole-cell system. Thus, we constructed NdmD mutants to explore the potential tunability of these enzymes.

Cells harboring wild-type NdmA or NdmB with either of the two reductase mutants showed a slight decrease in activity, although not nearly as drastic as that observed from purified proteins (Figure 5). The *N*_1_-demethylase activity in NdmA cells containing NdmDW and NdmDR was about 7% and 30% lower, respectively, than in cells with the wild type reductase. Conversely, *N*_3_-demethylase activity in NdmB cells was reduced by 36% and 20%, respectively, by cells with the NdmDW and NdmDR mutants. Thus, while these mutations do appear to interfere with the interaction between NdmD and either NdmA or NdmB, based on reduction of *N*-demethylase activity, the remaining activity is still more than 60% of the wild-type enzyme in our *in vivo* assays, demonstrating remarkable improvement in activity when compared with *in vitro* reactions.

**Figure 5.**
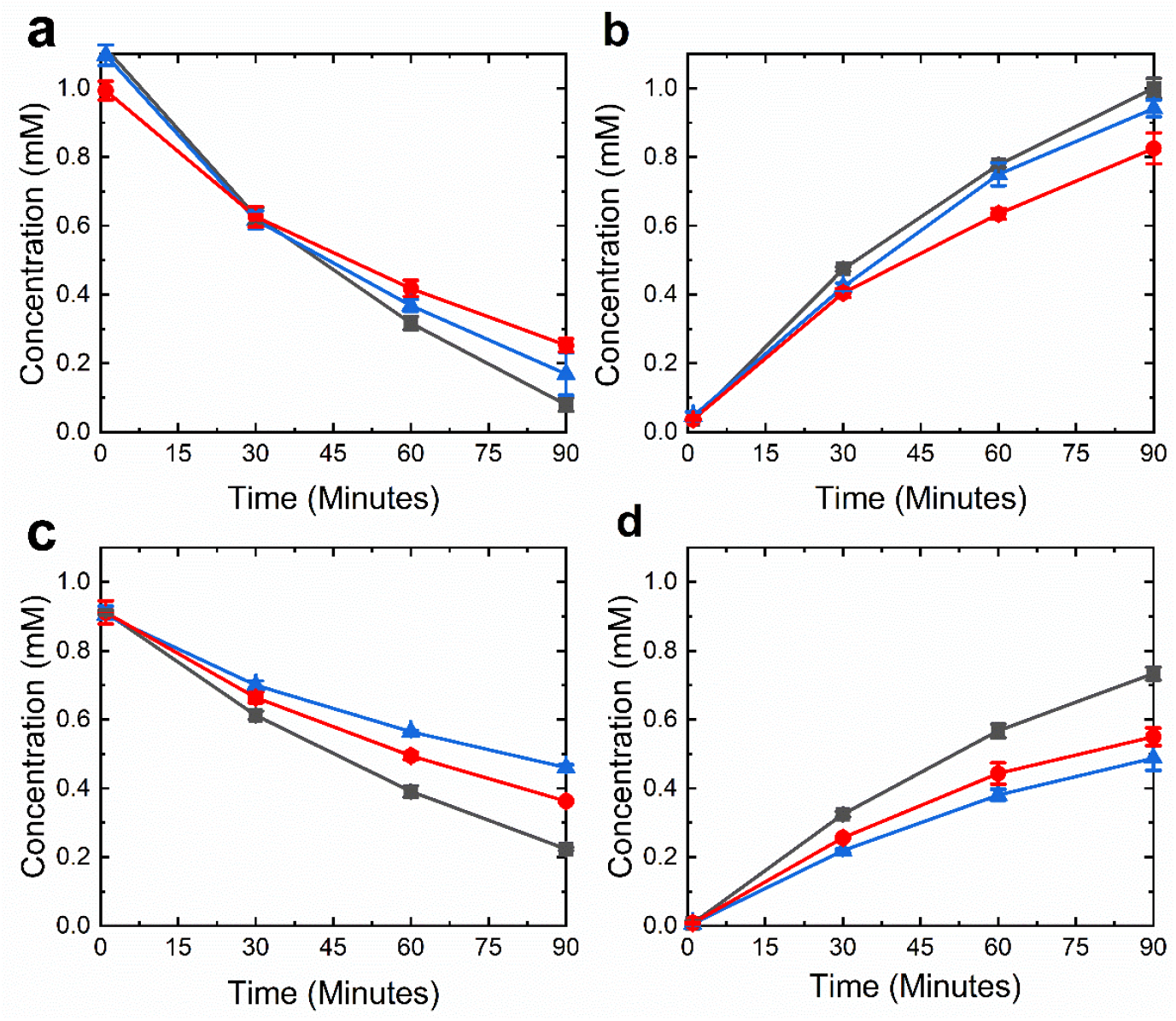
Whole cells containing NdmD, NdmDW, and NdmDR with NdmA or NdmB show a decrease in *N*-demethylation activity. Caffeine (a) is converted by NdmA to theobromine (b), and theobromine (c) is converted by NdmB to 7-methylxanthine (d). ■, NdmD; ▲, NdmDW; ●, NdmDR. Cells (OD_600_ = 5.0) were incubated with 1 mM caffeine or theobromine in 50 mM KP_i_ buffer at 30°C with 200 rpm shaking, and metabolites were quantified by HPLC. Concentrations reported are means with standard deviations of triplicate results.

The NdmD reductase contains an extra Rieske [2Fe-2S] cluster at its N-terminal end when compared with other Rieske reductases (Summers et al. 2013), but the cluster is not necessary for activity of NdmA (Retnadhas and Gummadi 2018). Thus, we also removed the Rieske cluster from the full-length NdmD by site-directed mutagenesis and assessed the effect of the mutations on *N*-demethylase activity of the whole-cell biocatalysts. Because the C50 and C69 residues of NdmD coordinate the Rieske [2Fe-2S] cluster, we mutated the residues to alanine so that the protein could not bind the cluster. First, a C69A single mutation was carried out on NdmD, NdmDW and NdmDR, resulting in NdmD1, NdmDW1, and NdmDR1, respectively. These single mutants were further altered to include a C69A mutation creating NdmD2, NdmDW2, and NdmDR2 double mutants.

Interestingly, we observed a wide range of activities for the various reductase mutants when they were assayed in our whole-cell biocatalyst system (Figures 6 and S2). There was no significant change in *N*_1_-demethylase activity of cells containing NdmA when NdmD was swapped for NdmD1 or NdmD2 (Figure 6A). This was as we expected given previous reports of a truncated reductase with similar activity (Retnadhas and Gummadi 2018). However, the single C69A and double C50A C69A mutations had a great effect on activity of the NdmDW and NdmDR mutants (Figure 6B). The rate of *N*_1_-demethylation was reduced by 18% with NdmDR1 and 58% by NdmDW1, when compared with NdmD1. The double mutation had a much greater effect on activity, as activity was reduced by 89% and 98% by NdmDR2 and NdmDW2, respectively, when compared with NdmD2.

**Figure 6.**
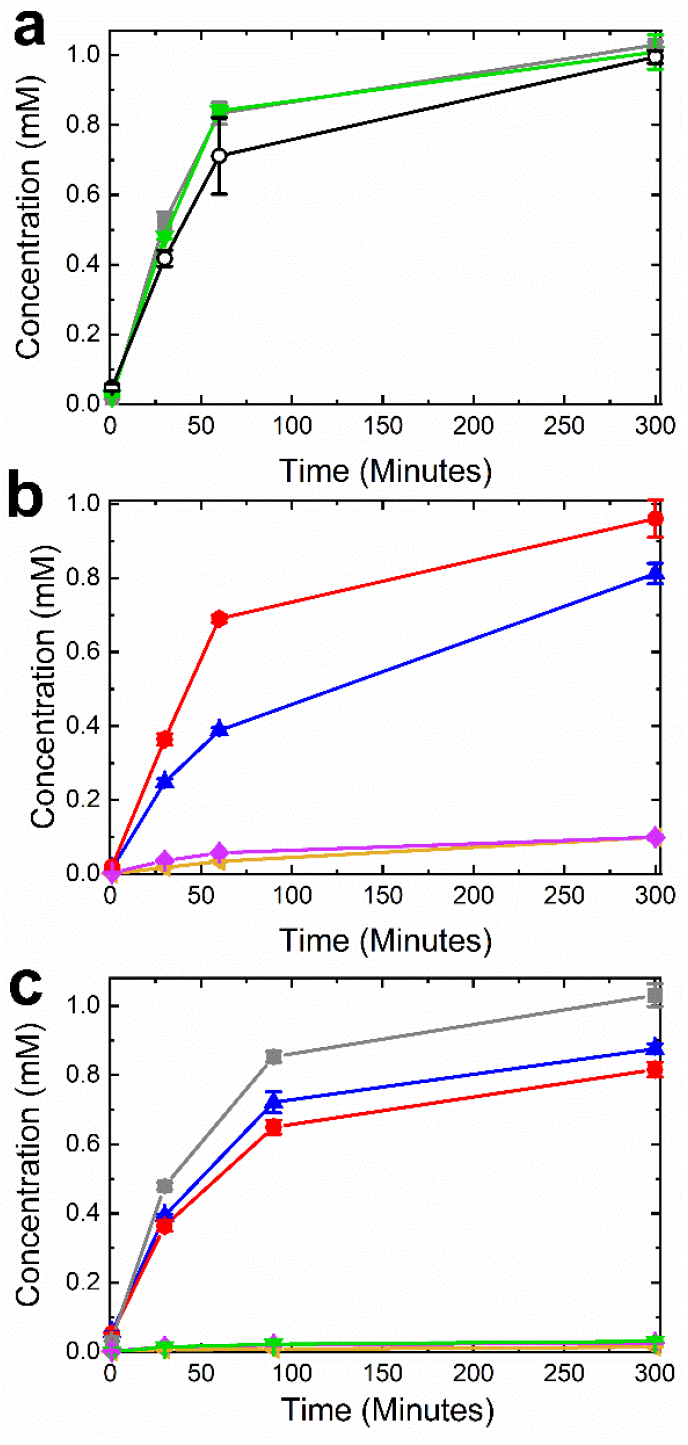
Mutations to the Rieske [2Fe-2S] binding site in NdmD result in various levels of *N*-demethylation activity. (a and b) Production of theobromine by cells containing NdmA, (c) production of 7-methylxnthine by cells containing NdmB. **○**, NdmD; ■, NdmD1; ●, NdmD2; ▲, NdmDW1; ◀, NdmDW2; ●, NdmDR1; and ♦, NdmDR2. Disappearance of substrate is shown in Figure S2. Concentrations reported are means with standard deviations of triplicate results.

For cells with NdmB, we also observed a range in *N*_3_-demethylation activity from the various mutant reductases assayed (Figure 6C). Compared with NdmD1, the activity of strains with NdmDR1 and NdmDW1 was reduced 33% and 27%, respectively. The activity of NdmD2, NdmDR2, and NdmDW2 mutants toward theobromine was almost negligible, with production of only 26.4 ± 2.0, 14.2 ± 0.4, and 30.8 ± 0.6 μM 7-methylxanthine after 5 hours.

## Discussion

Kim *et al*. established that the specificity of the *N-*demethylase enzymes was determined by the distance between the *N-*methyl group and the non-heme iron catalytic center (Kim et al. 2019). In order to change which *N-*methyl group is removed by NdmA or NdmB, site-directed mutagenesis was performed to alter the substrate binding pocket on the enzyme to try and rotate the caffeine substrate. The mutations to NdmB altered the rates of reaction with theobromine negatively, but were unable to change the enzyme selectivity from the *N*_3_-methyl group to the *N*_1_-methyl group on caffeine or theophylline. This is consistent with previous observations that the NdmB binding pocket may not be large enough to accommodate molecules with both *N*_1_- and *N*_3_-methyl groups (Summers et al. 2012).

In contrast to the NdmB results, mutations to NdmA did change the *N*-methyl positional selectivity of the enzyme toward caffeine and theophylline. The N282Q F286L double mutant, NdmA3, greatly shifted selectivity toward the *N*_3_-methyl group, and catalytic activity was further improved by swapping out a loop near the active site for the counterpart from NdmB. Resting cell assays with NdmA3 and NdmA4 cells accumulated significant amounts of paraxanthine from caffeine, which may enable biocatalytic production of paraxanthine in the future. Previous crystal structures of the NdmA3 enzyme with caffeine demonstrated that caffeine was bound in the reverse orientation as in NdmA, resulting in increased removal of the *N*_3_-methyl group (Kim et al. 2019). This report provides additional support for those findings and demonstrates this activity *in vivo*. The reduction in activity of NdmA4 toward theophylline compared with caffeine is curious. It may be that the β-loop from NdmB that was exchanged for the existing NdmA loop to create NdmA4 might interact with the *N*_7_-methyl group, stabilizing or guiding the substrate to the pocket. Yoneyama *et al*. determined that more than one amino acid was responsible for the recognition of purine derivatives and substrate discrimination in the theobromine synthase from *Camellia ptilophylla* and caffeine synthase from *Camellia sinensis*, and that the substrate specificity for xanthine derivatives is established by the residues in the central part of the enzyme (Yoneyama et al. 2006). Additionally, sequence alignment of the central 173 amino acids of each synthase revealed that there were only 9 amino acids that differed (Yoneyama et al. 2006). This is supported by the fact that the single mutations did not change the substrate positioning, only the rate of reaction. Our findings presented here align with this knowledge. While single mutations affected the reaction rates of both NdmA and NdmB, only the N282Q mutation on NdmA resulted in a large change in positional specificity. The theobromine and caffeine synthase enzymes essentially carry out the reverse reaction (*N*-methylation) compared with the *N*-demethylases in this study, but each must bind and properly align the methylxanthine for activity to occur.

Most metabolic engineering studies focus on modifying metabolism by altering gene expression, but recent studies have demonstrated that point mutations in enzymes can also be used to control flux of a compound through a metabolic pathway (Wu et al. 2020; Tovilla-Coutiño et al. 2020). The ability of mutations to either increase or decrease the overall reaction rate of catalytic reductases has been studied with some success in other enzymes. For example, error-prone PCR has been used in *Saccharomyces cerevisiae* to improve the cofactor binding of a xylose reductase, which resulted in an increase in production of nearly 40 times (Runquist et al. 2010). Additionally, distal point mutations within dihydrofolate reductase have been used to reduce the reaction rate and act as a probe to explore the relationship between the chemistry that the enzyme catalyzes and the protein structure beyond just the active site (Rajagopalan et al. 2002; Watney et al. 2003; Wang et al. 2006). Here, we have explored the potential of simple point mutations in the electron transfer protein NdmD to reduce the reaction rate of the *N*-demethylases NdmA and NdmB, creating several reductase mutants that lead to a wide range of *N*-demethylase activity in whole cell biocatalysts. This allows for fine-tuning of the overall process by controlling substrate conversion at the enzymatic level. Additional biochemical studies on purified mutant enzymes will be necessary to determine exactly how the mutations described here affect the electron transfer between the reductase and *N*-demethylase.

The data presented here are different from the mutant activity reported previously (Kim et al. 2019). In the previous report, the double mutant (NdmA3) exhibited slight activity toward caffeine, but activity of the loop-swapped double mutant (NdmA4) and reductase mutants (NdmDW and NdmDR) was almost negligible when using purified enzymes. By using whole-cell biocatalysts, we have demonstrated that NdmA4 does, indeed, demonstrate activity toward caffeine and that the reductase mutants show varying levels of activity *in vivo*. One explanation for the increased activity is the amount of enzyme present. We used a strong T7 promoter to drive expression of the *N*-demethylase genes, but have not accounted for solubility differences between the different mutant enzymes. Thus, our results are not meant to represent activity of a specific enzyme, but the activity of the whole-cell biocatalysts containing the desired enzymes.

Although we have generated new biocatalysts for production of methylxanthines, the catalytic rates could be improved with additional engineering. In order to optimize a binding site, there are three things to consider: the hydrogen bonding and van DerWaals interactions, the shape of the site relative to the substrate or ligand, and finally the structure of the protein in the unbound state to minimize entropy losses (Boehr et al. 2009; Fleishman et al. 2011). In addition to hydrogen bonds, polar interactions and ion pairing also regulate binding specificities (Schneider 1991; Dill 1990). Kim *et al*. identified the residues that interact directly with caffeine in the NdmA and NdmB binding pockets are due to hydrophobic interactions, and that the NdmA3 protein reverses the orientation of caffeine in the binding pocket (Kim et al. 2019). A computational model of the mutant enzyme would be beneficial to further optimize the hydrogen bonding and charges of the substrate and protein residues in order to increase enzyme solubility and activity. Hydrogen bonding specifically has been identified as one way to increase affinity of ligands or substrate with a mutant receptor, and optimization of charges in both binding pocket and adjacent residues can have an impact on specificity (Koh 2002). Mutations of residues near the binding or catalytic site are prone to affect substrate choice and offer new catalytic activities, enantioselectivity, and specificity (Dalby 2003; Park et al. 2005; Strausberg et al. 2005; Morley and Kazlauskas 2005). Other enzymatic properties may also be improved by mutations further out from the active site (Morley and Kazlauskas 2005).

While generation and screening of large mutant libraries for increased catalytic activity have great potential, in this case there are currently challenges in our ability to effectively screen mutants for increased production of a specific methylxanthine. The only current process to detect and quantify caffeine metabolites is by HPLC. Although this method is the most accurate way to monitor and detect caffeine degradation (Dash and Gummadi 2006), it is very time and labor intensive, and requires a large amount of materials. Development of new colorimetric or fluorescence-based methods to determine presence of specific caffeine metabolites will further increase the rate at which we can generate mutants for enhanced biocatalytic production of methylxanthines.

## Conclusion

In this work, we have determined the effect of distinct mutations on activity of the caffeine *N*_1_-demethylase NdmA expressed in whole cell biocatalysts and successfully swapped the specificity of NdmA from the *N*_1_- to the *N*_3_-methyl group of caffeine and theophylline. We have also demonstrated the potential to specifically control the rate of reaction using mutants of the NdmD reductase. Using this understanding, we have developed novel bacterial systems to expand our ability to produce high-value methylxanthine compounds, such as 1-methylxanthine and paraxanthine. Further optimization of the enzymes and strains can only improve upon the activity and stability shown here.

## Acknowledgements

We thank Dr. Hyun-Kyu Song of Korea University for generously sharing the pETDuet-NdmA4 plasmid. We thank Annabel Large and Madeline Stewart for their support and work in the lab on this project.

## Funding

This study was funded by University of Alabama research funds. M.B. Mock is supported by the U.S. Department of Education as a GAANN Fellow (P200A180056).

## Conflict of Interest

The authors declare no conflict of interest.

## Notes

### Competing Interest Statement

The authors have declared no competing interest.

