## Supplementary Information for "Rational Protein Engineering of Bacterial *N*-demethylases to Create Biocatalysts for the Production of Methylxanthines"

**Table S1.** All plasmids created and tested in this study.

| Plasmid Name | Plasmid Characteristics | Source |
|---|---|---|
| pACYCDuet-1 | P15A origin, two MCS, hexahistidine tag sequence by MCS1, Chloramphenicol resistance | Novagen |
| pETDuet-1 | pBR322 origin, two MCS, hexahistidine tag sequence by MCS1, Ampicillin resistance | Novagen |
| dDA | pACYCDuet-1 with NdmD and NdmA | (Algharrawi et al. 2015) |
| dDB | pACYCDuet-1 with NdmD and NdmB | (Algharrawi and Subramanian 2020) |
| dAA | pACYCDuet-1 with two copies of NdmA | (Algharrawi et al. 2015) |
| dBB | pACYCDuet-1 with two copies of NdmB | (Algharrawi and Subramanian 2020) |
| dDA1 | pACYCDuet-1 with NdmD and with NdmA mutation N282Q | This work |
| dDA2 | pACYCDuet-1 with NdmD and with NdmA mutation F286L | This work |
| dDA3 | pACYCDuet-1 with NdmD and with NdmA mutations N282Q and F286L | This work |
| dDB1 | pACYCDuet-1 with NdmD and with NdmB mutation L293F | This work |
| dDB2 | pACYCDuet-1 with NdmD and with NdmB mutation Q289N | This work |
| dDB3 | pACYCDuet-1 with NdmD and with NdmB mutations L293F and A289N | This work |
| pETDuet-NdmA4 | pETDuet-1 with NdmA4 | (Kim et al. 2019) |
| dDA4 | pACYCDuet-1 with NdmD and NdmA4 | This work |
| pET-28 a (+) | pBR322 origin, Kanamycin resistance | Novagen |
| pD | pET-28 a (+) with NdmD (previously named pET28-His-ndmD) | (Summers et al. 2012) |
| pD1 | pET-28 a (+) with NdmD mutation C69A | This work |
| pD2 | pET-28 a (+) with NdmD mutations C69A and C50A | This work |
| pDW1 | pET-28 a (+) with NdmD mutations C69A and V541W | This work |
| pDW2 | pET-28 a (+) with NdmD mutations C69A, C50A and V541W | This work |
| pDR1 | pET-28 a (+) with NdmD mutations C69A and V541R | This work |
| pDR2 | pET-28 a (+) with NdmD mutations C69A, C50A and V541R | This work |

**Table S2.** All primer sequences used in the plasmid construction for this study

| Primer Name | Primer Sequence 5′ to 3′ |
|---|---|
| A-N282Q-F | GATTATCTGCACATTGCATTTCAAGATCTCGTCTTCGCTGAAGAC |
| A-N282Q-R | GTCTTCAGCGAAGACGAGATCTTGAAATGCAATGTGCAGATAATC |
| A-F286L-F | GCATTTAATGATCTCGTCTTGGCTGAAGACAAACCAGTAATTG |
| A-F286L-R | CAATTACTGGTTTGTCTTCAGCCAAGACGAGATCATTAAATGC |
| A3-N282Q-F | GATTATCTGCACATTGCATTTCAAGATCTCGTCTTGGCTGAAGAC |
| A3-N282Q-R | GTCTTCAGCCAAGACGAGATCTTGAAATGCAATGTGCAGATAATC |
| B-L293F-F | GCTTTCCAGAAGCGGGTGTTTGACGAAGACCAGCCTG |
| B-L293F-R | CAGGCTGGTCTTCGTCAAACACCCGCTTCTGGAAAGC |
| B-Q289N-F | CACATGCACCTGGCTTTCAACAAGCGGGTGCTTGACGAAG |
| B-Q289N-R | CTTCGTCAAGCACCCGCTTGTTGAAAGCCAGGTGCATGTG |
| B3-Q298N-F1 | CACATGCACCTGGCTTTCAACAAGCGGGTGTTTGACGAAG |
| B3-Q289N-R1 | CTTCGTCAAACACCCGCTTGTTGAAAGCCAGGTGCATGTG |
| D-V541W-F | GAAGCTTCTTGTGAGCAGGGTTGGTGCGGGACTTGTATAACTCCAG |
| D-V541W-R | CTGGAGTTATACAAGTCCCGCACCAACCCTGCTCACAAGAAGCTTC |
| D-V541R-F | GAAGCTTCTTGTGAGCAGGGTCGCTGCGGGACTTGTATAACTCCAG |
| D-V541R-R | CTGGAGTTATACAAGTCCCGCAGCGACCCTGCTCACAAGAAGCTTC |
| D-C50A-F | AATGCTTGGGAGAACCGCGCCCCGCATAGAGGATTGCGG |
| D-C50A-R | CCGCAATCCTCTATGCGGGGCGCGGTTCTCCCAAGCATT |
| D-C69A-F | GCTAATACCGGTAACGAGTTGCGAGCTCAGTATCATGGATGGACTTATG |
| D-C69A-R | CATAAGTCCATCCATGATACTGAGCTCGCAACTCGTTACCGGTATTAGC |
| Loop-F-NdeI | GCACGGCATATGGAACAGGCAATCATTAATG |
| NdmA-R-KpnI | CCTCCGGGTACCTTATARGTAGCTCCTATCGCTT |

**Table S3.** All strains created and tested in this study.

| **Strain Name** | **Strain Characteristics** | **Source** |
|---|---|---|
| *E. coli* BL21(DE3) | $F^-$ *ompT hsdS*$_B$ ($r_B^- m_B^-$) *gal dcm* (DE3) | Invitrogen |
| *E. coli* dDA | BL21(DE3) dDA | (Algharrawi et al. 2015) |
| *E. coli* dDA1 | BL21(DE3) dDA1 | This work |
| *E. coli* dDA2 | BL21(DE3) dDA2 | This work |
| *E. coli* dDA3 | BL21(DE3) dDA3 | This work |
| *E. coli* dDA4 | BL21(DE3) dDA4 | This work |
| *E. coli* dDB | BL21(DE3) dDB | (Algharrawi and Subramanian 2020) |
| *E. coli* dDB1 | BL21(DE3) dDB1 | This work |
| *E. coli* dDB2 | BL21(DE3) dDB2 | This work |
| *E. coli* dDB3 | BL21(DE3) dDB3 | This work |
| *E. coli* pDdAA | BL21(DE3) pD dAA | (Algharrawi et al. 2015) |
| *E. coli* pD1dA | BL21(DE3) pD1 dAA | This work |
| *E. coli* pD2dAA | BL21(DE3) pD2 dAA | This work |
| *E. coli* pDWdAA | BL21(DE3) pDW dAA | This work |
| *E. coli* pDW1dAA | BL21(DE3) pDW1 dAA | This work |
| *E. coli* pDW2dAA | BL21(DE3) pDW2 dAA | This work |
| *E. coli* pDRdAA | BL21(DE3) pDR dAA | This work |
| *E. coli* pDR1dAA | BL21(DE3) pDR1 dAA | This work |
| *E. coli* pDR2dAA | BL21(DE3) pDR2 dAA | This work |
| *E. coli* pD1dBB | BL21(DE3) pD1 dBB | This work |
| *E. coli* pD2dBB | BL21(DE3) pD2 dBB | This work |
| *E. coli* pDWdBB | BL21(DE3) pDW dBB | This work |
| *E. coli* pDW1dBB | BL21(DE3) pDW1 dBB | This work |
| *E. coli* pDW2dBB | BL21(DE3) pDW2 dBB | This work |
| *E. coli* pDRdBB | BL21(DE3) pDR dBB | This work |
| *E. coli* pDR1dBB | BL21(DE3) pDR1 dBB | This work |
| *E. coli* pDR2dBB | BL21(DE3) pDR2 dBB | This work |

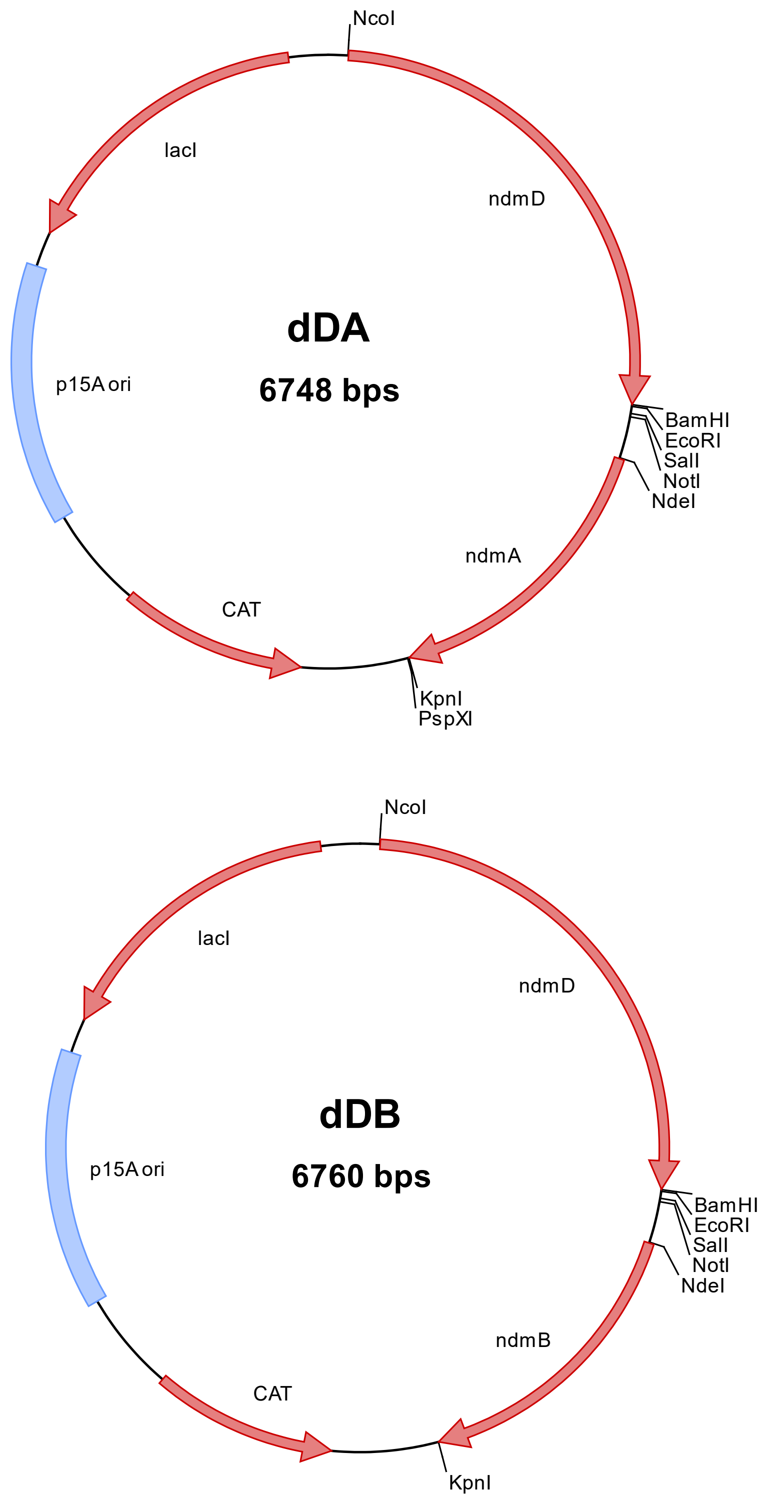


**Fig. S1** Plasmid maps of dDA and dDB, which served as the basis for all further plasmids created in this work.

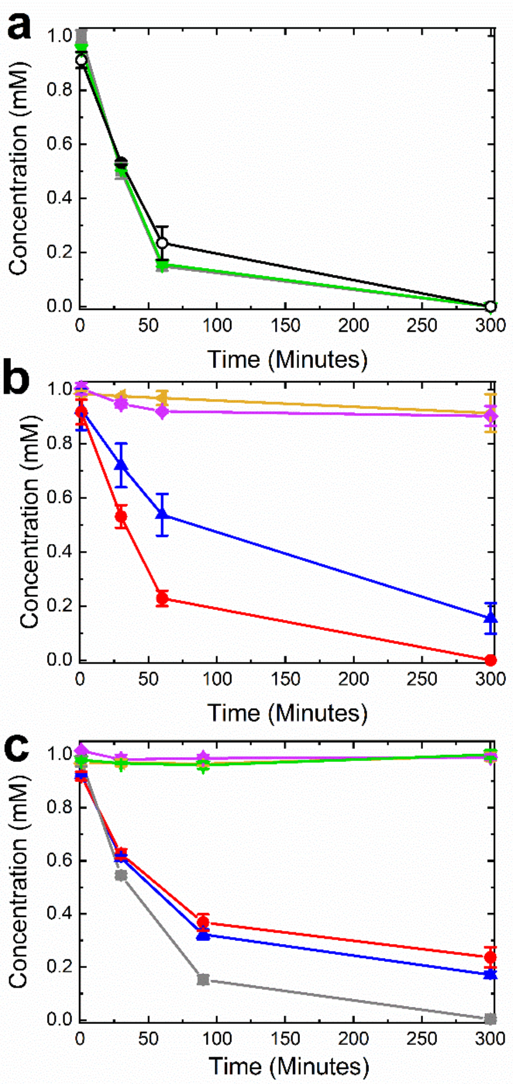


**Fig. S2** Substrate conversion by whole cells expressing *ndmA* or *ndmB* with various *ndmD* mutants show varying levels of activity. Caffeine is converted to theobromine by cells expressing *ndmA* with (a) ○, *ndmD*, ■, *ndmD1* and ●, *ndmD2*, or (b) ▲, *ndmDW1*, ◄, *ndmDW2*, ●, *ndmDR1*, and ♦, *ndmDR2*. (c) Theobromine is converted to 7-methylxanthine by cells expressing *ndmB* with ■, *ndmD1*, ▲, *ndmDW1*, ●, *ndmDR1*, ▼, *ndmD2*, ◄, *ndmDW2* and ♦, *ndmDR2*. Cells ($OD_{600}$ = 5.0) were incubated with 1 mM caffeine or theobromine in 50 mM $KP_i$ buffer at 30°C with 200 rpm shaking, and metabolites were quantified by HPLC. Concentrations reported are means with standard deviations of triplicate results.
